# Genome assembly reconstruction of the Japanese honey bee, *Apis cerana japonica* (Hymenoptera: Apidae), using homology-based assembly and nanopore long-reads

**DOI:** 10.1101/2023.07.26.550500

**Authors:** Yudai Masuoka, Akiya Jouraku, Seigo Kuwazaki, Mikio Yoshiyama, Mari Horigane-Ogihara, Taro Maeda, Yutaka Suzuki, Hidemasa Bono, Kiyoshi Kimura, Kakeru Yokoi

## Abstract

Honey bees are important for agriculture (e.g., pollination and honey production). Additionally, honey bees are an important insect model species, especially as model social insects. The Japanese honey bee, *Apis cerana japonica* (a subspecies of the Asian honey bee, *Apis cerana*), is a Japanese domestic honey bee, which has several subspecies-specific traits. We previously constructed the draft genome sequence data of *A. cerana japonica*, but it needed to be improved considering the use of the genome sequence data for genome structural analysis and repetitive region analysis, as well as the availability of chromosome-level genome data of *A. mellifera* and *A. cerana*. In this study, we constructed the improved *A. cerana japonica* genome data and new gene set data with functional annotations. The constructed genome data, including 16 pseudochromosomes, was found to be highly contiguous and complete, and the gene set data covered most of the core genes in the BUSCO database. Thus, the constructed genome and gene set data have become more suitable as the reference data of *A. cerana japonica*.

## 1. Introduction

Honey bees are important for pollination, honey production, and as model social insect species^1^. Among the *Apis* species, the Western honey bee, *Apis mellifera*, is used worldwide in apiculture because *A. mellifera* has good traits for humans, such as their highly producing honey compared to the other wild *Apis* species. Considering its importance, the genome of *A. mellifera* was sequenced and reported in 2006^2^, and the genome and gene set data have been improved over time^3,4^. Now, the chromosome-level assembly data and gene set data of *A. mellifera* based on this data are available. These data have helped advance the research using *A. mellifera* at the genetic and molecular levels^5^. Furthermore, multiple genome sequences of *A. mellifera* subspecies, including chromosome-level genome sequence data, have been published and are currently available in the public genome database^6,7^. The genome data of other *Apis* species have also been reported^8,9^. *A. cerana*, an Asian honey bee, has several different traits compared with *A. mellifera*, such as resistance to Varroa mites^10^ (summarized by Park et al., 2015^11^). The first genome sequence data of *A. cerana*, which is a Korean native, was reported in 2015^11^, and the genome sequence data of other *A. cerana* from China was published in 2018^12^. Subsequently, chromosome-level genome sequence data of the native Chinese species were reported (hereafter, referred to as *A. cerana cerana*)^13^. *A. cerana japonica* (a subspecies of *A. cerana*) is the Japanese domestic honey bee and has several subspecies-specific traits (e.g., hot defensive bee ball against Vespa^14,15^). To clarify the traits at the molecular or genetic level and promote comparative research using the genome data of *Apis* species, we previously reported the draft genome sequence data of *A. cerana japonica*^16^. The size of the draft *A. cerana japonica* genome data is 211 Mbp, N50 is 180 kbp, and the number of predicted genes is 13,222, which are comparable to those of the other *Apis* species.

Considering the use of *A. cerana japonica* genome data for genome structural analysis and repetitive region analysis, including transposable elements (TE)^6^, satellite DNA, and tandem duplications (such analysis was performed in *Drosophila* species^17^), and considering that more contiguous sequence data are available for *A. mellifera* and *A. cerana*^3,13^, the draft genome data of *A. cerana japonica* needed to be improved. In this study, we updated the draft genome data of *A. cerana japonica*, and the improved genome sequence data was constructed. Specifically, we scaffolded the draft genome data using the high-quality genome sequence data of *A. cerana cerana* as a reference, and gap closing of the scaffolded genome data was performed using the newly sequenced *A. cerana japonica* long-read genome data. In addition, gene predictions were performed on the newly constructed genome data using RNA-Seq data as a hint. The analyses of the reconstructed genome and gene set data revealed that these data are more suitable as the reference data of *A. cerana japonica*, which can contribute to honey bee research at the genetic and molecular levels and support comparative genome research.

## 2. Materials and Methods

### 2.1 Sample preparation and sequencing

Genomic DNA (gDNA) of *A. cerana japonica* was extracted from a single male larva in the prepupal stage using the MagAttract HMW DNA kit (QIAGEN, Venlo, Nederland) according to the manufacturer’s protocol. MinION libraries were prepared from 1 _μ_g of long gDNA using the 1D squared Ligation Sequencing Kit (SQK-LSK308, Oxford Nanopore Technologies, Oxford, UK) according to the manufacturer’s protocol. Sequencing was performed using the MinION sequencer with a FLO-MIN107 flowcell for 72 h with the MinKNOW software. Raw sequence data of the gDNA was deposited in the DNA Data Bank of Japan (DDBJ) Sequence Read Archive (SRA) (Accession ID: DRR385865, BioSample ID: SAMD00509887, BioProject ID: PRJDB13846).

Total RNA was extracted for RNA-Seq from each whole body of a foraging adult worker and each whole body of workers, which are 7 to 16 days (larva to adult) after oviposition. Each whole body sample was homogenized in 1 mL of TRIzol Reagent (Thermo Fisher Scientific, Waltham, Massachusetts, U.S.A) using metal beads (5 mm). Each homogenate was incubated for 5 min at room temperature, and 0.2 mL of chloroform was added and mixed. Each sample was centrifuged at 14,000 × g for 10 min at 4°C. The upper, aqueous phase was transferred to a new tube. An equal volume of 70% ethanol was added and mixed using a vortex mixer. Each sample was transferred to an RNeasy Mini spin column and purified according to the RNeasy Mini Kit (QIAGEN) protocol. The extracted total RNA of foraging adults and larvae was sequenced using the Illumina HiSeq 4000 and Illumina NovaSeq 6000 systems, respectively (Macrogen Japan Corp., Tokyo, Japan). All RNA-Seq data were deposited in the SRA (Accession IDs and other related IDs are shown in Supplemental Table S1)

### 2.2 Genome data construction, searching for the repetitive region and transposable elements and assessments of the genome data and predicted gene set data

The previous draft genome assembly data of *A. cerana japonica* that we constructed (GenBank assembly accession ID: GCA_002217905.1)^16^ was used as an input genome assembly and improved using the following methods. First, using the chromosome-level genome assembly of *A. cerana cerana* (GenBank assembly accession ID: GCA_011100585.1)^13^ as a reference, the homology-based scaffolding was performed for the input genome assembly using the scaffold command in RagTag software (version 2.1.0)^18^. Second, the high-quality nanopore-long reads were filtered using the NanoFilt software (version 2.8.0)^19^, and the filtered sequence data was applied to close gaps in the new genome assembly using TGS-GapCloser software (version 1.2.0)^20^. The sequence data of the constructed *A. cerana japonica* genome assembly were deposited in DDBJ/ENA/GenBank (GenBank accession IDs: BSDG01000001-01000240, Supplemental Table S2). For searching repetitive regions and TEs, a de novo repeat library of consensus sequences of TEs for the new genome assembly was constructed using RepeatModeler2 (version 2.0.1)^21^, and repetitive regions and TEs in the new genome assembly were searched in the sequence data using RepeatMasker (version 4.1.0)^22^ and the constructed repeat library. The RepeatMasker analysis was performed using Dfam_3.1 and rmblastn version 2.10.0+, in which repetitive regions and TEs were masked for gene prediction input data. Whole genome alignment with *A. cerana cerana* was performed using LASTZ software (version 1.04.15)^23^. For the validation of the constructed genome sequence data and predicted gene set data, including the predicted amino acid sequence data, other *Apis* genera, including *A. mellifera* (GenBank assembly accession ID: GCA_003254395.2)^3^ and *A. cerana cerana* genome data, and the previous draft genome data were analyzed for comparison using the BUSCO software (version 5.3.2) with the Insecta_Odb10 dataset (2020-09-10) in gVolante (URL: https://gvolante.riken.jp/)^24,25^.

### 2.3 Gene prediction and functional annotation

The RNA-Seq raw reads were trimmed using fastp software (version 0.22.0)^26^ and mapped on the new genome assembly using STAR software (version 2.7.10a)^27^ to make a BAM file. Subsequently, a gene set on the new genome assembly was predicted using AUGUSTUS software (version 3.3.3)^28^ with hint data generated from the BAM file using masked sequence data as the input. Gene annotations of the predicted amino acid sequences were performed using Fanflow4Insects^29^ with data of eight reference proteins from model organisms and insect species (*A. mellifera, Homo sapiens, Mus musculus, Caenorhabditis elegans, D. melanogaster, Bombyx mori, Bombus terrestris*, and *Nasonia vitripennis*) or Unigene protein sequence data with the default settings.

## 3 Results and Discussion

### 3.1. Construction of *Apis cerana japonica* genome assembly

A new genome assembly of *A. cerana japonica* was constructed by homology-based scaffolding using the previous draft genome assembly and the *A. cerana cerana* chromosome-level genome assembly as a reference^13^, and gap closing of the constructed genome data was performed using long-read sequences obtained by nanopore sequencing. The number of scaffolds, average length, maximum length of scaffolds, and N50 value of the new genome assembly are 240, approximately 905 kbp, 27.49 Mbp, and 12.86 Mbp, respectively (Table 1). These values are an improvement over the draft genome data of *A. cerana japonica* and comparable to the other *Apis* genome data at the chromosome level. Moreover, the total length and GC content of the new assembly is approximately 217 Mbp and 32.79%, which is similar to those of other *Apis* genome data (Table 1). These results suggested that the quality and continuity of the *A. cerana japonica* genome data were significantly improved from the draft genome data. However, the median and minimum lengths of the genome data were not improved because the genome data contains short-size sequence data, which were not scaffolded. The 240 contigs contained 16 pseudochromosomes and 224 unplaced scaffolds. The total length of the 16 pseudochromosomes was approximately 208.68 Mb, covering 95.99% of the total length of the whole genome. The genomic synteny comparison with the 16 chromosomes of *A. cerana cerana*, which was used as a reference, showed highly conserved synteny, as expected (Fig. 1). Therefore, these contigs reflected chromosomes, and the contiguous genome assembly that was constructed can be useful in the analyses of gene structures, regulatory regions of expression, repetitive regions and TEs, as performed previously using the genome data of *Drosophila* species^17^.

**Table 1.** Basic statistics of genome assembly in the *Apis* genus.

|  | *A. cerana japonica*<br>(Yokoi et al., 2018)<sup>16</sup> | *A. cerana japonica*<br>(this study) | *A. cerana cerana*<br>(Wang et al., 2020)<sup>13</sup> | *A. mellifera*<br>(Wallberg et al., 2019)<sup>3</sup> |
| --- | --- | --- | --- | --- |
| Total length (bp) | 211,196,132 | 217,405,261 | 215,670,033 | 225,250,884 |
| Number of sequences | 3,315 | 240 | 126 | 177 |
| Average lengths (bp) | 63,710 | 905,855 | 1,711,667 | 1,272,604 |
| Median lengths (bp) | 22,323 | 4799.5 | 25,330.5 | 19,344 |
| Sequence size with maximum length (bp) | 1,316,057 | 27,491,524 | 26,804,424 | 27,754,200 |
| Sequence size with minimum length (bp) | 242 | 242 | 3,587 | 2,303 |
| N50 | 180,259 | 12,866,345 | 12,422,783 | 12,619,445 |
| GC content (%) | 33.04 | 32.79 | 32.79 | 33 |
| Number of genes | 13,222 | 12,000 | 10,741 | 9,935 |

**Fig 1.**
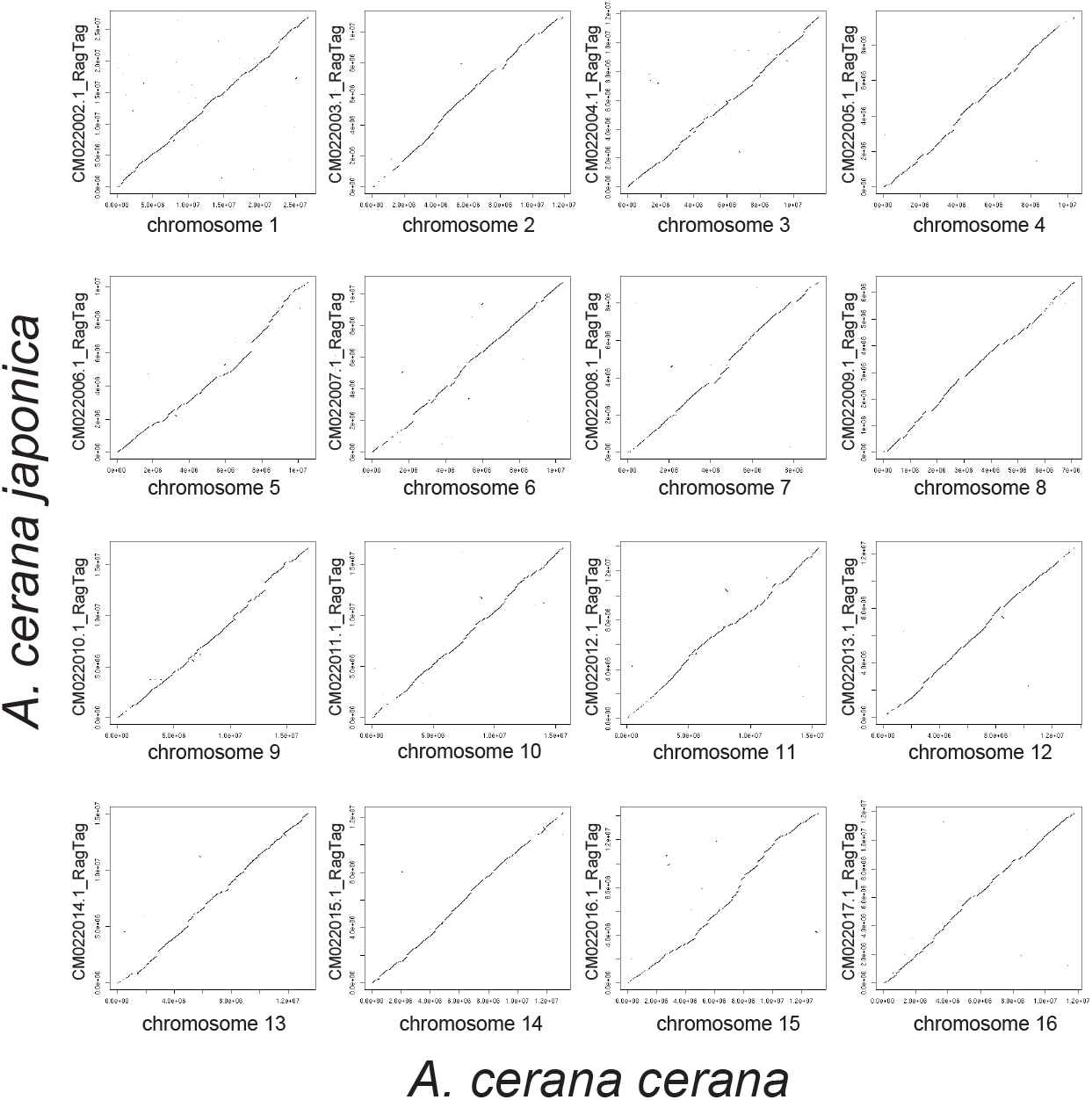
Whole genome alignment between *Apis cerana japonica* and *Apis cerana cerana*. Notably, 16 pseudochromosomes of *A. cerana japonica* were constructed and aligned with *A. cerana cerana*. All 16 pseudochromosomes in *A. cerana japonica* showed highly conserved synteny with *A. cerana cerana* chromosomes. The ID on the left side of each graph indicates the contig ID of the gff file corresponding to each reference chromosome.

### 3.2 Detections of repetitive regions and transposable elements

Repetitive regions and TEs in the new genome assembly were searched using RepeatModeler2 and RepeatMasker (the summary of the results and the output data of RepeatModeler2 and RepeatMasker were in Table 2 and Supplemental data S1, respectively), showing that 8.80% of the assembly consists of repetitive regions or TEs, which is comparable to other *Apis* species^31^. The TEs detected in the assembly comprised retroelements (Class I), including LINEs and LTR elements, and the DNA transposons (Class II) included Tc1-IS630-Pogo, which is consistent with other *Apis* species and the previous *A. cerana japonica* genome^6^. Furthermore, the number of the annotated TEs per category was counted, as performed in our previous reports, for comparison with other *Apis* species and the previous draft genome data^6^ (Table 3). Three Class I TEs were detected, namely LINE/R1, LINE/R2, and LTR/Copia, and more of these Class I TEs (LINE/R1: 113, LINE/R2: 95, and LTR/Copia: 658) were detected than those in the draft genome (LINE/R1: 57, LINE/R2: 51, and LTR/Copia: 82), whereas LTR/ERVK and LTR/Gypsy, which were found in the draft genome, were not detected in the updated genome assembly. Additionally, the results showed that the detected Class II TEs consisted of DNA/CMC-EnSpm, DNA/Dada (not detected in the draft genome), DNA/TcMar-Mariner, and DNA/TcMar-Tc1, whereas DNA/hAT-Ac, which was detected in the draft genome, was not detected in the updated genome assembly. Additionally, more TEs of DNA/TcMar-Mariner and DNA/TcMar-Tc1 (DNA/TcMar-Mariner: 852 and DNA/TcMar-Tc1: 306) were detected in the updated assembly data than in the draft genome, and less TEs of DNA/CMC-EnSpm were detected in the updated assembly data than in the draft genome (Table 3). Since many TEs have been junked in the process of evolution (fragmentations or mutations of their sequences) and since some TEs share common characteristics, it is conceivable that slight differences in the parameters of the application being used could result in different TE classifications. DNA/Data was newly detected, and less TEs of DNA/CMC-EnSpm were detected in the current assembly. However, more TEs of LINE/R1, LINE/R2, LTR/Copia, DNA/TcMar-Mariner, and DNA/TcMar-Tc1 may have been detected in the updated assembly because they are located in the genome regions, of which the sequences in the draft genome data were not determined.

**Table 2.**
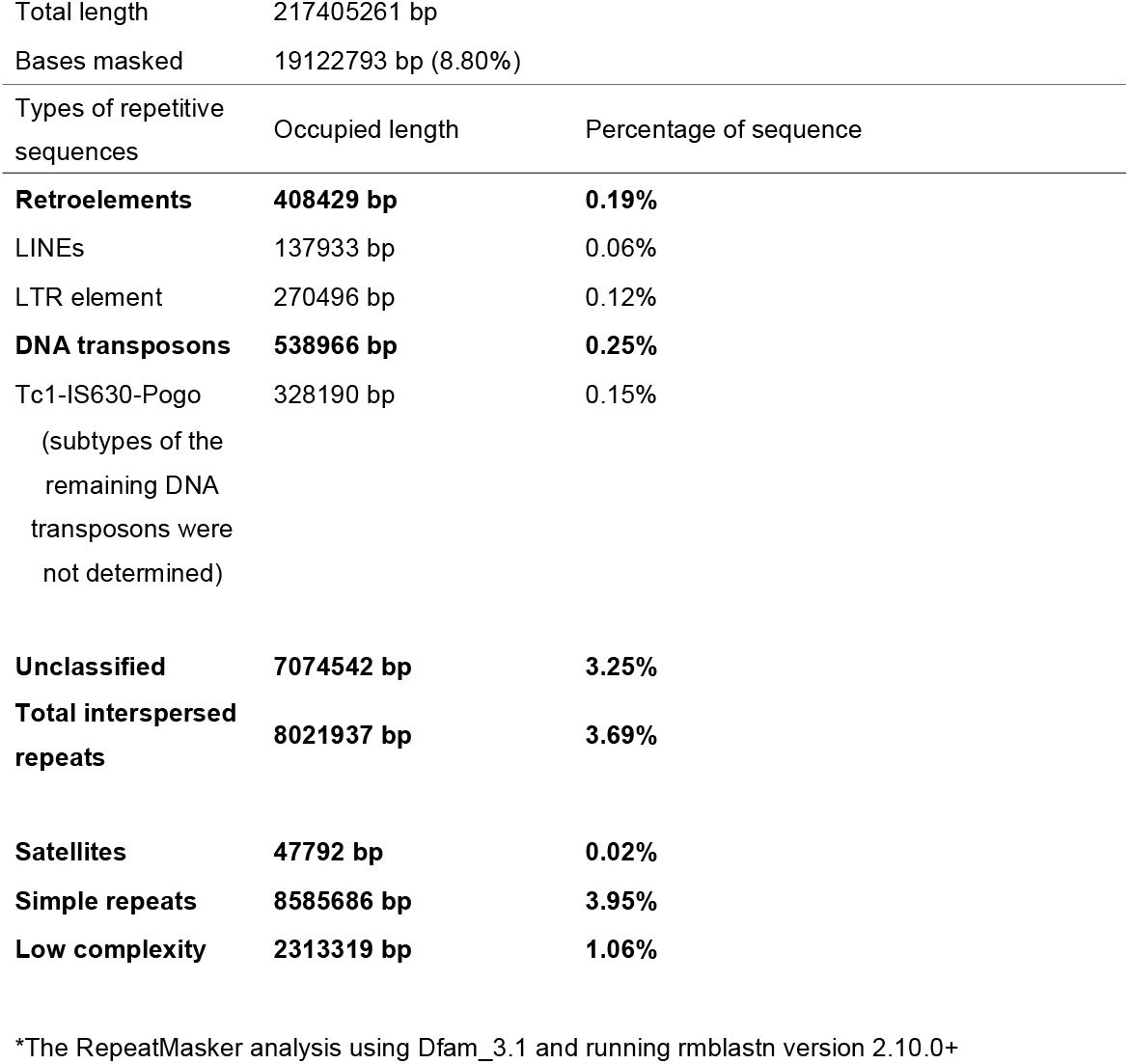
Summary of repetitive element analysis using RepeatMasker.

| Total length | 217405261 bp |  |
| --- | --- | --- |
| Bases masked | 19122793 bp (8.80%) |  |
| Types of repetitive sequences | Occupied length | Percentage of sequence |
| **Retroelements** | **408429 bp** | **0.19%** |
| LINEs | 137933 bp | 0.06% |
| LTR element | 270496 bp | 0.12% |
| **DNA transposons** | **538966 bp** | **0.25%** |
| Tc1-IS630-Pogo<br>(subtypes of the<br>remaining DNA<br>transposons were<br>not determined) | 328190 bp | 0.15% |
| **Unclassified** | **7074542 bp** | **3.25%** |
| **Total interspersed repeats** | **8021937 bp** | **3.69%** |
| **Satellites** | **47792 bp** | **0.02%** |
| **Simple repeats** | **8585686 bp** | **3.95%** |
| **Low complexity** | **2313319 bp** | **1.06%** |

\*The RepeatMasker analysis using Dfam\_3.1 and running rmbblastn version 2.10.0+

**Table 3.** Numbers of TEs annotated in the new assembly data. Each number in the parenteses indicates each TE number detected in the draft genome^6^.

| Class I |  |  | Class II |  |  |  |
| --- | --- | --- | --- | --- | --- | --- |
| LINE/R1 | LINE/R2 | LTR/Copia | DNA/CMC-EnSpm | DNA/Dada | DNA/TcMar-Mariner | DNA/Tc |
| 113 (57) | 95 (51) | 658 (82) | 588 (1684) | 616 (0) | 852 (630) | 306 |

### 3.3 Gene prediction and functional annotation of predicted genes

The repetitive regions and sequence regions annotated as TE in the constructed genome data were masked using RepeatMasker for gene predictions. Using the masked genome data and RNA-Seq data of *A. cerana japonica* which were mapped to the new genome assembly as hint data, 12,000 genes were predicted (Supplemental data S2), which is comparable to the number of predicted genes for *A. cerana cerana* and *A. mellifera* (Table 1)^3,13^. For assessing the completeness of the assembled genomes, BUSCO software was used. Although the percentage of the “complete” category (98.7%) for the new genome assembly is slightly lower than that of *A. cerana cerana* and *A. mellifera* (99.4%), most BUSCO core genes were found in the updated genome data, and the percentage of the “complete” category is higher than that of the previous draft genome data (97.6%) (Table 4). Furthermore, the BUSCO results obtained using the new predicted amino acid data revealed that the predicted gene set data covered most of the BUSCO core genes in the Insecta_Odb10 dataset (97.5%). These results suggested that our reconstructed genome data and predicted gene set data possess high completeness, and they are more suitable than the previous draft data for detailed comparative analysis of genome sequences (e.g., genome structure analysis) or gene sets with other species or subspecies such as *A. mellifera* and *A. cerana cerana* at the chromosome level^3,13^.

**Table 4.** The results of BUSCO using the constructed genome sequence data.

|  | *A. cerana japonica*<br>(Yokoi et al., 2018) | *A. cerana japonica*<br>(This study) | *A. cerana cerana*<br>(Wang et al.,<br>2020) | *A. mellifera*<br>(Wallberg et al.,<br>2019) | *A. cerana*<br>*japonica*<br>peptide (This<br>study) |
| --- | --- | --- | --- | --- | --- |
| Complete | 97.6% (1334/1367) | 98.7% (1349/1367) | 99.4%<br>(1359/1367) | 99.4% (1359/1367) | 97.5%<br>(1348/1367) |
| Single-copy | 96.6% | 95.1% | 99.3% | 99.3% | 94.1% |
| Duplicated | 1.0% | 3.6% | 0.1% | 0.1% | 3.4% |
| Fragmented | 1.1% | 0.4% | 0.1% | 0% | 1.0% |
| Missing | 1.3% | 0.9% | 0.5% | 0.6% | 1.5% |

The predicted genes were functionally annotated using Fanflow4Insects (Supplemental data S3)^29^. Using individual amino acid sequences, Fanflow4Insects performed homologous searches against multiple model and insect species, and hmmscan with the Pfam dataset. Table 5 summarizes the number of hits and hit ratio of the amino acid sequences against each database. Although less than 60% of the predicted amino acid data were hits against the *C. elegans* dataset, approximately 70% to 80% of the amino acid data were hits against the other dataset, except for Pfam. Such high percentages suggested that the annotation results can be used for biological interpretations in omics analyses (e.g., enrichment analysis by meta-scape^30^). However, approximately 20% of the remaining genes were not functionally annotated. Some of these genes may be involved in *Apis-*specific or *Apis cerana japonica*-specific traits.

**Table 5.**
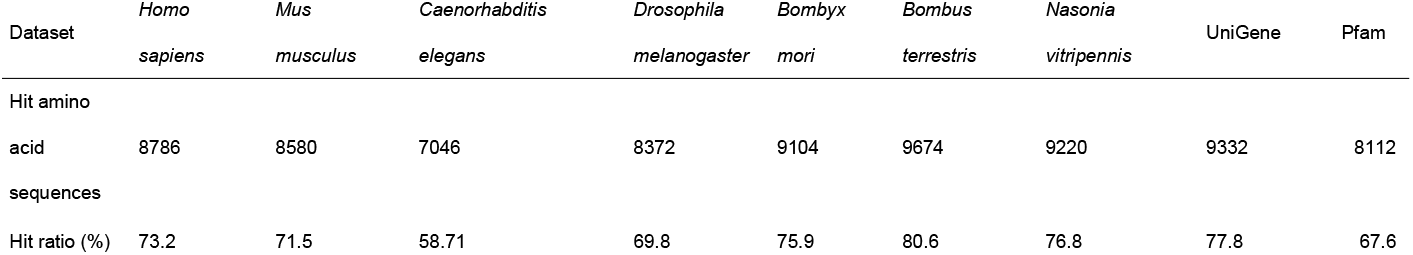
Numbers of annotated genes of *A. cerana japonica* against reference protein data of multiple model species and insects, Unigene or Pfam domain scan.

## 4. Conclusion

In this study, a high-quality genome assembly, including 16 pseudochromosomes, was constructed by scaffolding using the *A. cerana cerana* chromosome-level genome assembly as a reference and by gap closing using nanopore long-read sequences. The basic status of the constructed genome data, such as N50 value, total length, GC content ratio of the constructed genome assembly, as well as the results of synteny and BUSCO analysis, suggested that the draft data was sufficiently improved and may serve as a useful reference for *A. cerana japonica*. Furthermore, the gene set data containing 12,000 genes with functional annotations were constructed using the masked genome data and RNA-Seq data. BUSCO analysis showed that the gene set data possessed high completeness, covering most of BUSCO core genes and possessing sufficient quality as a reference, which can be used for future research and comparative genome analysis between *Apis* species or others.

## Acknowledgments

We are grateful to Masatsugu Hatakeyama for productive discussions and help in writing manuscripts.

We dedicate this paper to Dr. Panuwan Chantawannakul. She passed away spring of 2022 and the world is less without her warm presence. She was interested in the genome study of Apis, and was the inspiration to start this Japanese honey bee genome analysis project.

## Supplemental Materials

All supplemental materials are available in figshare. DOI: 10.6084/m9.figshare.c.6455737

### Supplemental Table S1

Accession numbers of raw RNA-Seq data for *Apis cerana japonica* gene predictions. DOI:10.6084/m9.figshare.22210963.

### Supplemental Table S2

Accession IDs of the assembled genome sequences (IDs in gff file, raw description of fasta file, and DDBJ accession IDs). DOI: 10.6084/m9.figshare.22220404.

### Supplemental data S1

Output files of RepeatModeler2 and RepeatMasker. The fasta (new_Acj.fa) and stk (new_Acj.stk) files contain consensus sequences of TEs in *Apis cerana japonica* from RepeatModeler2. The two files, new_Acj.out and new_Acj.tbl, are the result files from RepeatMasker. new_Acj.masked.gz is the genome sequence file of *Apis cerana japonica*, in which repetitive regions and TE sequences are shown in lower cases (used input file for gene prediction). DOI: 10.6084/m9.figshare.22220488.

### Supplemental data S2

Output files of AUGUSTUS. “new_Acj_gene.gtf” shows structural information of the predicted genes in the assembled genome.“new_Acj_gene_cds.fa” and “new_Acj_gene_aa.fa” are fasta files containing the protein-coding sequences and translated amino acid sequences, respectively. DOI: 10.6084/m9.figshare.22221004.

### Supplemental data S3

Functional annotation results of the predicted genes using fanflow4Insects^29^. DOI: 10.6084/m9.figshare.22221106.

